# Enhancing mitochondrial pyruvate metabolism ameliorates myocardial ischemic reperfusion injury

**DOI:** 10.1101/2024.02.01.577463

**Authors:** Joseph R. Visker, Ahmad A. Cluntun, Jesse N. Velasco-Silva, David R. Eberhardt, Thirupura S. Shankar, Rana Hamouche, Jing Ling, Hyoin Kwak, Yanni Hillas, Ian Aist, Eleni Tseliou, Sutip Navankasattusas, Dipayan Chaudhuri, Gregory S. Ducker, Stavros G. Drakos, Jared Rutter

## Abstract

The established clinical therapy for the treatment of acute myocardial infarction is primary percutaneous coronary intervention (PPCI) to restore blood flow to the ischemic myocardium. PPCI is effective at reperfusing the ischemic myocardium, however the rapid re-introduction of oxygenated blood also can cause ischemia-reperfusion (I/R) injury. Reperfusion injury is the culprit for up to half of the final myocardial damage, but there are no clinical interventions to reduce I/R injury. We previously demonstrated that inhibiting the lactate exporter, monocarboxylate transporter 4 (MCT4), and re-directing pyruvate towards oxidation can blunt isoproterenol-induced hypertrophy. Based on this finding, we hypothesized that the same pathway might be important during I/R. Here, we establish that the pyruvate-lactate metabolic axis plays a critical role in determining myocardial salvage following injury. Post-I/R injury, the mitochondrial pyruvate carrier (MPC), required for pyruvate oxidation, is upregulated in the surviving myocardium following I/R injury. MPC loss in cardiomyocytes caused more cell death with less myocardial salvage, which was associated with an upregulation of MCT4 in the myocardium at risk of injury. We deployed a pharmacological strategy of MCT4 inhibition with a highly selective compound (VB124) at the time of reperfusion. This strategy normalized reactive oxygen species (ROS), mitochondrial membrane potential (Δψ), and Ca^2+^, increased pyruvate entry to TCA cycle, and improved myocardial salvage and functional outcomes following I/R injury. Altogether, our data suggest that normalizing the pyruvate-lactate metabolic axis via MCT4 inhibition is a promising pharmacological strategy to mitigate I/R injury.

**GRAPHICAL ABSTRACT:** 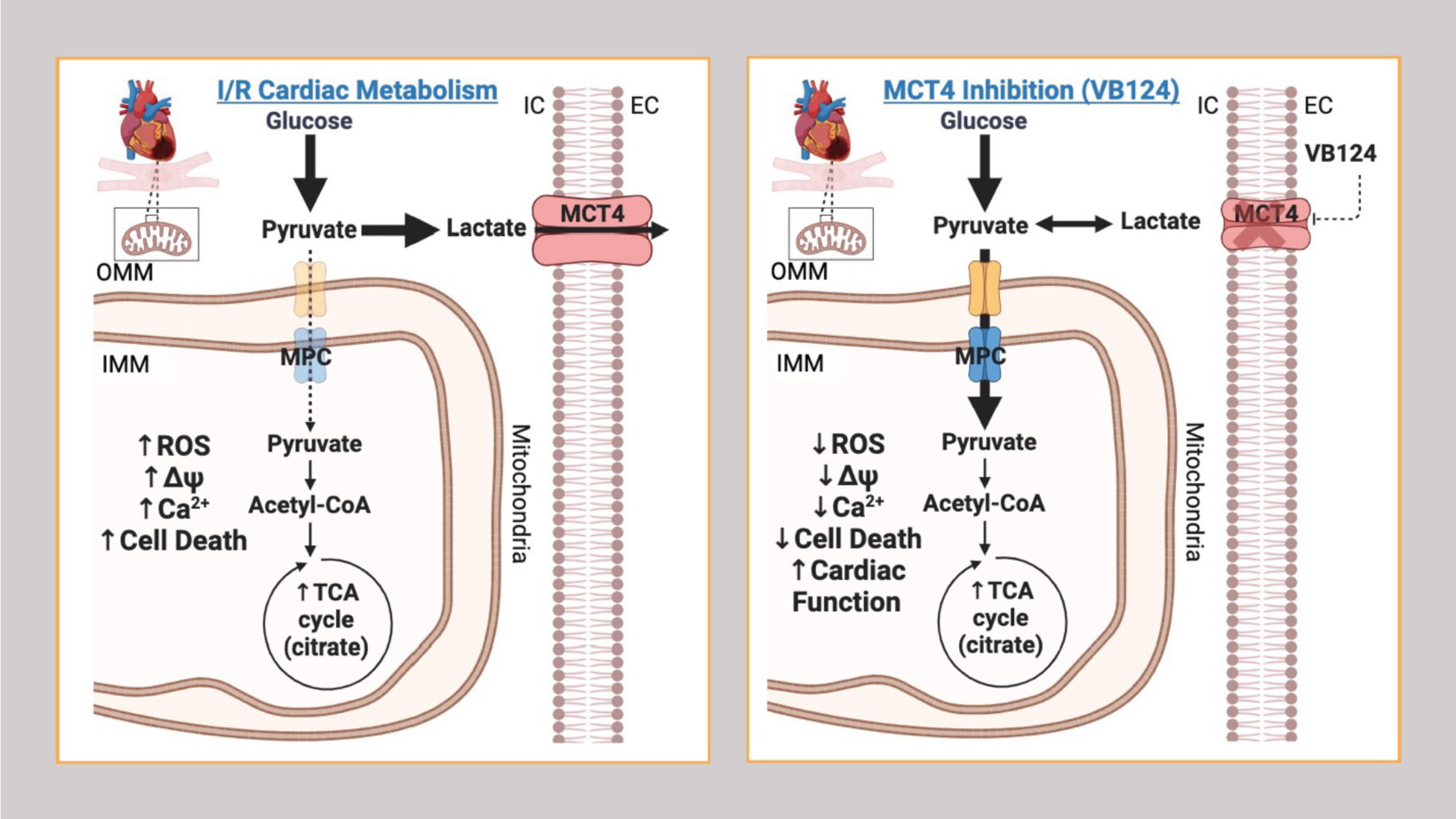

## INTRODUCTION

Acute myocardial infarction (AMI) is a major public health problem and a principal cause of heart failure (HF), which is a leading cause of mortality worldwide ^1–3^. The standard of care for AMI patients is primary percutaneous coronary intervention (PPCI) to reperfuse and restore oxygenated blood flow to the ischemic myocardium ^4, 5^. Paradoxically, PPCI is accompanied by reperfusion injury, which exacerbates tissue injury and increases cardiomyocyte death resulting in a reduction of salvageable myocardium. It is estimated that reperfusion injury accounts for ∼50% of the final infarct after an AMI ^4,6^. Despite decades of research, there are no pharmacological interventions that have been successfully translated into routine clinical practice to alleviate the detrimental effects of I/R injury ^7–9^ . Therefore, mitigation of myocardial I/R injury remains an unmet need in cardiovascular medicine to prevent the development of chronic HF following ischemic events.

The mechanisms underlying I/R are complex and multifactorial, but data from animal models suggests that a key contributor is mitochondrial dysfunction within the ischemic cardiomyocytes ^10–12^. Mitochondrial function is critical in cardiomyocytes during I/R injury to maintain cellular energetics, redox and viability ^13^. Mitochondrial defects in response to I/R injury, which can result in mitochondria-mediated apoptosis, include impaired mitochondrial membrane potential, calcium overload and oxidative stress ^14, 15^. This is thought to arise in I/R due to a metabolic imbalance caused by the discontinuous availability of oxygen and nutrients during ischemia and reperfusion ^16, 17^. Understanding the metabolic perturbations associated with I/R could enable the development of novel cardioprotective therapies to minimize the damage associated with I/R and prevent subsequent HF.

The mitochondrial pyruvate carrier (MPC) is an inner mitochondrial membrane obligate heterodimeric transporter (composed of MPC1 and MPC2), which facilitates entry of pyruvate into the mitochondrial matrix where it can be converted into acetyl-CoA to feed the tricarboxylic (TCA) cycle ^18^. In the relatively hypoxic fetal heart, cardiac energy production is met largely by carbohydrate fuels and glycolytic metabolism. This metabolism also enables biomass production for the growing heart. However, a metabolic transition occurs at birth and the mature myocardium derives ∼60-90% of its ATP from fatty acid oxidation ^19^.

During an ischemic event, the supply of oxygen and nutrients to the myocardium is reduced, which is partially compensated by an increase in glucose consumption, a build-up of electrons in the form of NADH and succinate, and increased production of lactate due to the disruption in aerobic respiration ^20, 21^. Upon reperfusion of the ischemic myocardium, the sudden influx of oxygen and nutrients elicits a metabolic imbalance that causes a surge of reactive oxygen species (ROS) resulting from rapid succinate oxidation ^11, 12^. The MPC is upregulated in the peri-infarct border zone of I/R injury and may be cardioprotective by contributing to increased energy production, reduced oxidative stress, and the promotion of pro-survival pathways ^22^. We and others recently showed that adult cardiomyocyte-specific deletion of MPC1 caused cardiac hypertrophy and HF ^23–25^. When pyruvate cannot enter the TCA cycle via the MPC it is converted into lactate by the activity of lactate dehydrogenase (LDH). Lactate is exported from cardiomyocytes by the activity of monocarboxylate transporter 4 (*SLC16A3* or MCT4) ^20, 21, 26^. We have defined this dynamic interplay between pyruvate and lactate as the “pyruvate-lactate metabolic axis”^18^.

Utilizing both *in vitro and in vivo* models of I/R, we now show that cardiomyocyte-specific deletion of MPC1 leads to less myocardial salvage and greater overall injury. Moreover, *MPC1*-deficient (*MPC1*^CKO^) hearts exhibited a ∼20% increase in myocardial necrosis following I/R, which was accompanied by increased MCT4 expression. Based on these observations, we tested the hypothesis that inhibiting lactate export and re-directing pyruvate carbon through the MPC into mitochondrial oxidation would be protective in models of I/R injury. We employed the small-molecule MCT4 inhibitor VB124 in *in vitro* and *in vivo* models of acute and chronic I/R injury ^18^. We found that MCT4 inhibition normalized ROS, mitochondrial membrane potential, Ca^2+^ levels, and pyruvate oxidation, leading to increased myocardial salvage and improved cardiac function. Therefore, building upon an understanding of the metabolic mechanisms surrounding I/R, our data suggest that targeting the pyruvate-lactate metabolic axis with MCT4 inhibition is a promising cardioprotective therapy to ameliorate I/R injury.

## RESULTS

### Murine hearts lacking the MPC have less myocardial salvage and more necrosis following I/R injury

We used a cardiac-specific, tamoxifen-inducible conditional *MPC1* deletion mouse model to investigate the role of MPC1 in the myocardial response to I/R (Figure 1A). At 4 weeks post injection (WPI) with tamoxifen, and prior to any observable cardiovascular phenotype, we subjected *MPC1^CKO^* mice (Figure 1B) to *in vivo* myocardial I/R injury (Figure 1C) and assessed the extent of myocardial damage. Using the double-staining histological technique of Evans blue and triphenyltetrazolium chloride (TTC) ^27–31^, we controlled for infarct size and found that the area at risk (AAR) was indistinguishable between WT and MPC1^CKO^ hearts, indicating a similar initial ischemic insult (Figure 1D). However, when normalized to the AAR, the myocardial salvage was reduced by 20% in MPC1^CKO^ mice compared to their paired WT littermates (Figure 1E-G).

**Figure 1:**
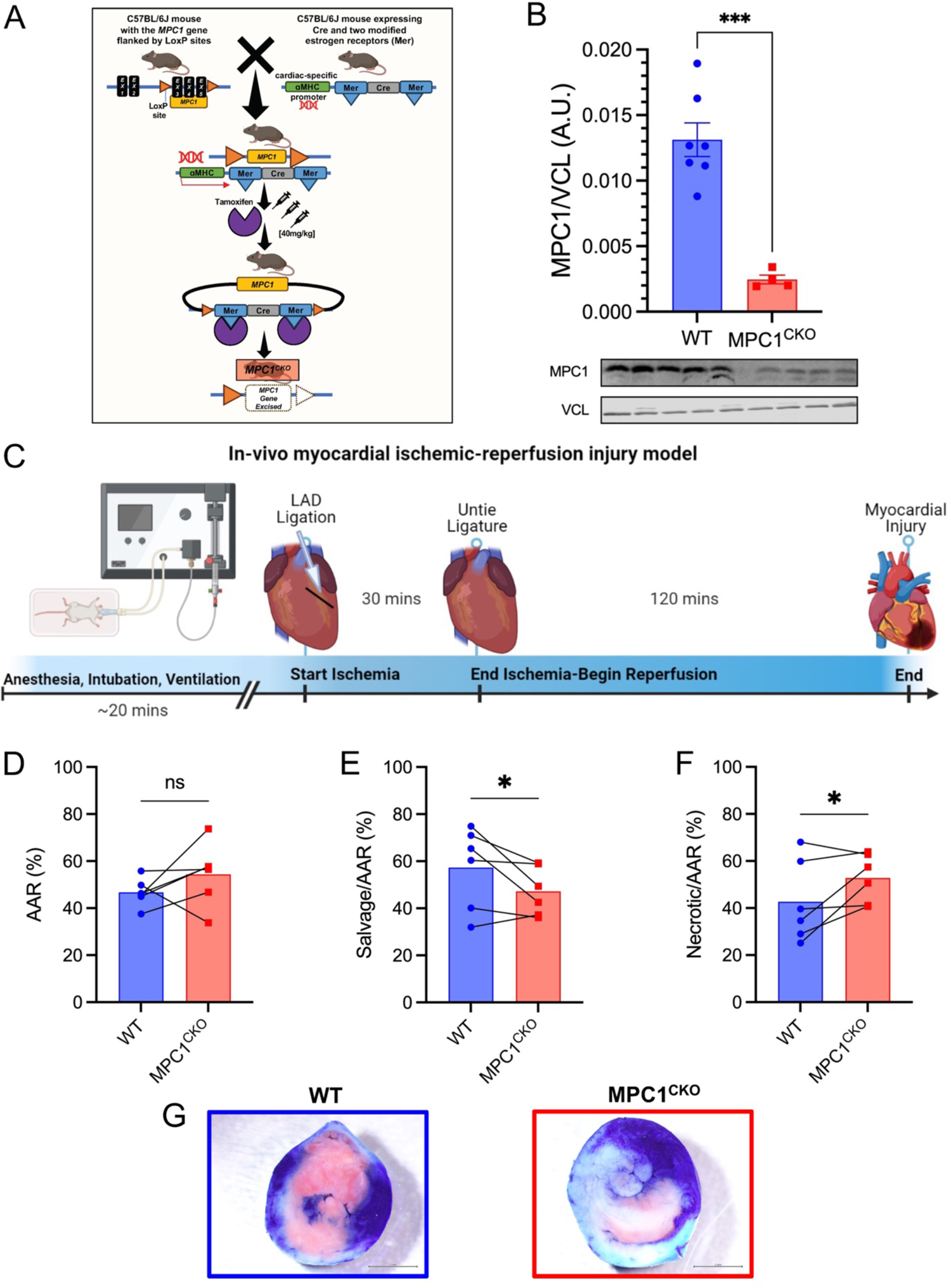
Loss of the MPC in murine hearts results in less myocardial salvage with more necrosis following I/R. **A)** The MPC1 gene locus is targeted in C57Bl/6J mice by placing loxP sites in the introns flanking exons 3-5 of the *MPC1* genomic locus. Cardiomyocyte specificity was engineered by crossing MPCfl/fl mice with an MHC driven tamoxifen inducible Cre. To induce deletion of *MPC1* (MPC1^CKO^), mice were intraperitoneally injected with tamoxifen for three consecutive days (40 mg/kg) at 8 weeks old. **B)** Western blot showing significant knockdown of MPC1 from whole hearts. **C)** Schematic of *in vivo* myocardial I/R injury model. **D)** Following I/R injury, the area at risk (AAR%) is non-significant (p=0.13), indicating similar initial ischemic injury between WT (46.67±2.45%) and MPC1^CKO^ (54.33±5.42%). **E & F)** Within the area at risk, myocardial salvage (TTC: pink tissue staining) is significantly reduced (p=0.04) in the MPC1^CKO^ (47.21±4.22%, n=6) when compared to their paired WT littermates (57.28±7.09%, n=6). Representative images of myocardial salvage, and necrosis following *in vivo* I/R injury in WT and MPC1^CKO^ mice. Paired t-test was used for statistical analysis. * = p<0.05, *** = p<0.001, ns=non-significant.

### MPC1^CKO^ mice upregulate several deleterious pathways upon I/R

To understand the underlying mechanisms associated with the differential response to I/R injury in the MPC1^CKO^ and WT mice, we performed quantitative transcriptomics on hearts. Again, we subjected both groups to I/R and following two hours of reperfusion, separated the heart into ischemic tissue (peri-infarct area), and non-ischemic tissue (away from the infarct) before RNA was isolated and sequenced (Figure 2A). To identify differentially expressed genes, we used a 5% false discovery rate with DeSeq2 software (version 1.24.0), and genes were filtered using an adjusted p-value <0.05, and absolute log2 fold change > 0.585. We performed ingenuity pathway analysis (IPA) as an integrative approach to identify enriched groups of genes among those that were differentially expressed.

**Figure 2:**
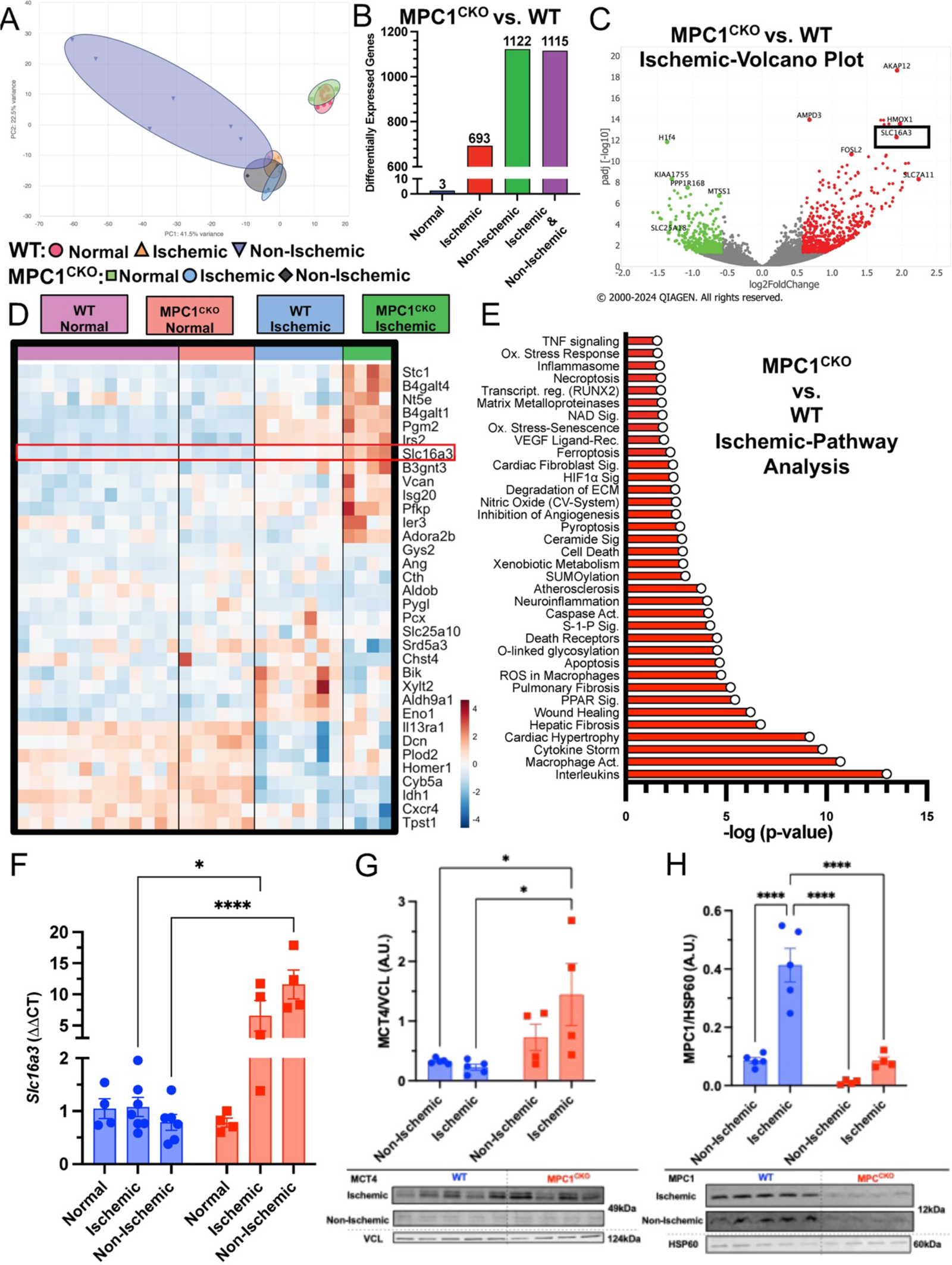
MPC1^CKO^ mice upregulate several deleterious pathways upon I/R. **A)** Principal component analysis plot showing gene expression profiles of WT (normal: n=13, ischemic: n=7, non-ischemic: n=7) and MPC1^CKO^ (normal: n=6, ischemic: n=4, non-ischemic: n=4) datasets. **B)** Bar graph showing the distribution of significant genes within experimental conditions. **C)** Volcano plot of genes identified using IPA in the MPC1^CKO^ ischemic myocardium, including *Slc16a3* (MCT4, black box). **D)** Heatmap of genes associated with glycolysis in WT and MPC1^CKO^ hearts and their response to I/R injury showing an upregulation of *Slc16a3* in MPC1^CKO^ hearts. **E)** IPA pathway analysis reveals that within the ischemic tissue of MPC1^CKO^ hearts there is an association with deleterious pathways such as interleukin signaling, macrophage activation, cytokine storm activity, and cell death. **F)** Quantitative real-time polymerase chain reactions (qRT-PCR) showing *Slc16a3* expression is normally low in both WT and MPC1^CKO^ hearts but upon I/R injury *Slc16a3* gene expression is upregulated in the ischemic and non-ischemic samples of only the MPC1^CKO^. **G and H)** The relative protein abundance via western blotting of MCT4 and MPC1 in the non-ischemic and ischemic myocardium reveal an unbalanced pyruvate-lactate metabolic axis following I/R injury. Unpaired t-tests and 2-way ANOVAs were used for statistical analysis between WT (n=5) and MPC1^CKO^ (n=4), and ischemic and non-ischemic tissue. * = p<0.05, ** = p<0.01, *** = p<0.001, ns=non-significant.

To test whether the transcriptional changes that occurred were due to I/R intervention, we also obtained RNA from healthy WT and MPC1^CKO^ hearts at 4 WPI, before MPC1^CKO^ hearts show any cardiovascular phenotype. As expected, I/R injury led to large changes in gene expression (Supplemental Figure 1). In response to I/R injury, there were 1,820 differentially expressed genes with a significant change (p<0.05) in the MPC1^CKO^ hearts compared to WT after multiple comparisons correction. Of these, 693 genes were differentially expressed in the ischemic myocardium, closest to the injury site (Figure 2B). Away from the myocardial injury, within the non-ischemic myocardium, there was 1,122 genes differentially expressed in the MPC1^CKO^ heart compared to WT (Supplemental Figure 2). Importantly, at this short time post-tamoxifen, there were almost no gene expression differences between WT and MPC1^CKO^ hearts.

The volcano plot in Figure 2C depicts the genes that are differentially expressed between ischemic WT and MPC1^CKO^ heart tissue. We highlight that the solute carrier family 16 member 3 (*Slc16a3:* red dot, block box), which encodes the lactate exporter MCT4, was the eighth most significantly upregulated gene in the ischemic myocardium of the MPC1^CKO^ tissue. This is consistent with the hypothesis that lactate production and excretion provide an alternative route of pyruvate disposal when mitochondrial import and oxidation are blocked due to the loss of the MPC. The heatmap in Figure 2D shows the differential expression of a broader set of genes associated with carbohydrate metabolism in response to I/R injury in the MPC1^CKO^ hearts.

IPA analysis revealed an upregulation of several deleterious pathways in MPC1^CKO^ ischemic myocardium compared to WT, such as interleukin signaling (p=9.45E^-14^), macrophage activation (p=1.79E^-11^), cytokine storm activity (p=1.59E^-10^), and cell death (p=1.48E^-05^), all of which may promote tissue injury following I/R (Figure 2E). We additionally examined gene signatures of cardiotoxic processes within MPC1^CKO^ hearts such as cardiac infarction (p=2.13E^-12^), fibrosis (p=1.02E^-11^), cell death (p=4.80E^-10^), and dysfunction (p=1.61E^-10^), compared to WT littermates (Supplemental Figure 3). In summary, following I/R, there is clear evidence of exacerbated tissue damage with a stress-associated transcriptional phenotype in MPC1^CKO^ hearts, including evidence of metabolic rewiring that includes upregulation of the *Slc16a3* gene.

### MPC1^CKO^ hearts express high levels of MCT4 upon I/R

We next performed quantitative real-time polymerase chain reaction (qRT-PCR) analysis and immunoblotting to quantify *Slc16a3* mRNA and MCT4 and MPC1 protein expression in ischemic and non-ischemic tissue of WT and MPC1^CKO^ mouse hearts. Without I/R, *Slc16a3* gene expression is low and similar in both groups. Upon I/R injury, however, *Slc16a3* expression significantly increases in the WT ischemic and non-ischemic heart tissue, which is exacerbated in the MPC1^CKO^ hearts (Figure 2F, qRT-PCR). To determine whether the *Slc16a3* gene expression pattern translates to increased protein abundance (Figure 2G), we performed western blotting and found that relative MCT4 protein abundance in MPC1^CKO^ mice was increased in the ischemic but not in the non-ischemic myocardial tissue when compared to WT.

Interestingly, we note that the observed changes in MCT4 protein abundance post-I/R are reciprocal to those of MPC1 (Figure 2H). MPC1 protein abundance within the ischemic area is significantly upregulated in WT mice, but as expected is much lower in the MPC1^CKO^ mice upon I/R. Our data show that at rest the myocardium lacking the MPC has low expression levels of *Slc16a3* and MCT4, similar to WT animals. Upon I/R injury, WT animals increase expression of MPC1 within the ischemic area, but in the MPC1^CKO^ heart MCT4 instead is induced. Whether this is adaptive or maladaptive is unclear.

### MPC1^CKO^ cardiomyocytes under simulated I/R exhibit greater mitochondrial damage, increased cell death, and more lactate/succinate flux

To understand if there exist mitochondrial defects in MPC1^CKO^ cardiomyocytes that may contribute to enhanced injury upon I/R, we utilized an *in vitro* model to simulate I/R. Following a previously described protocol, 4 WPI primary adult cardiomyocytes (ACMs) derived from WT and MPC1^CKO^ mice were isolated and exposed to two hours of hypoxia followed by two hours of reoxygenation (Figure 3A) ^32^. ACMs were then stained with mitochondria-specific reporters for ROS (MitoSOX), membrane potential (tetramethylrhodamine methyl ester, TMRM), or calcium (X-Rhod1) and imaged. MPC1^CKO^ ACMs cultured in normoxia showed less ROS, and membrane potential (Δψ) compared to WT ACMs at baseline, potentially suggesting a lower energetic state prior to I/R injury (Figure 3B-E). However, following simulated I/R injury, we observed increases in mitochondrial ROS, Δψ, and Ca^2+^ levels in both groups. The MPC1^CKO^ cells demonstrated greater fold increases in ROS and Δψ, but not Ca^2+^ (Figure 3B). Consistent with our *in vivo* phenotype, simulated I/R injury resulted in significantly more cell death in MPC1^CKO^ cells compared to controls (Figure 3F). Whether the measured differences in mitochondrial function can explain differences in survival remains unclear.

**Figure 3:**
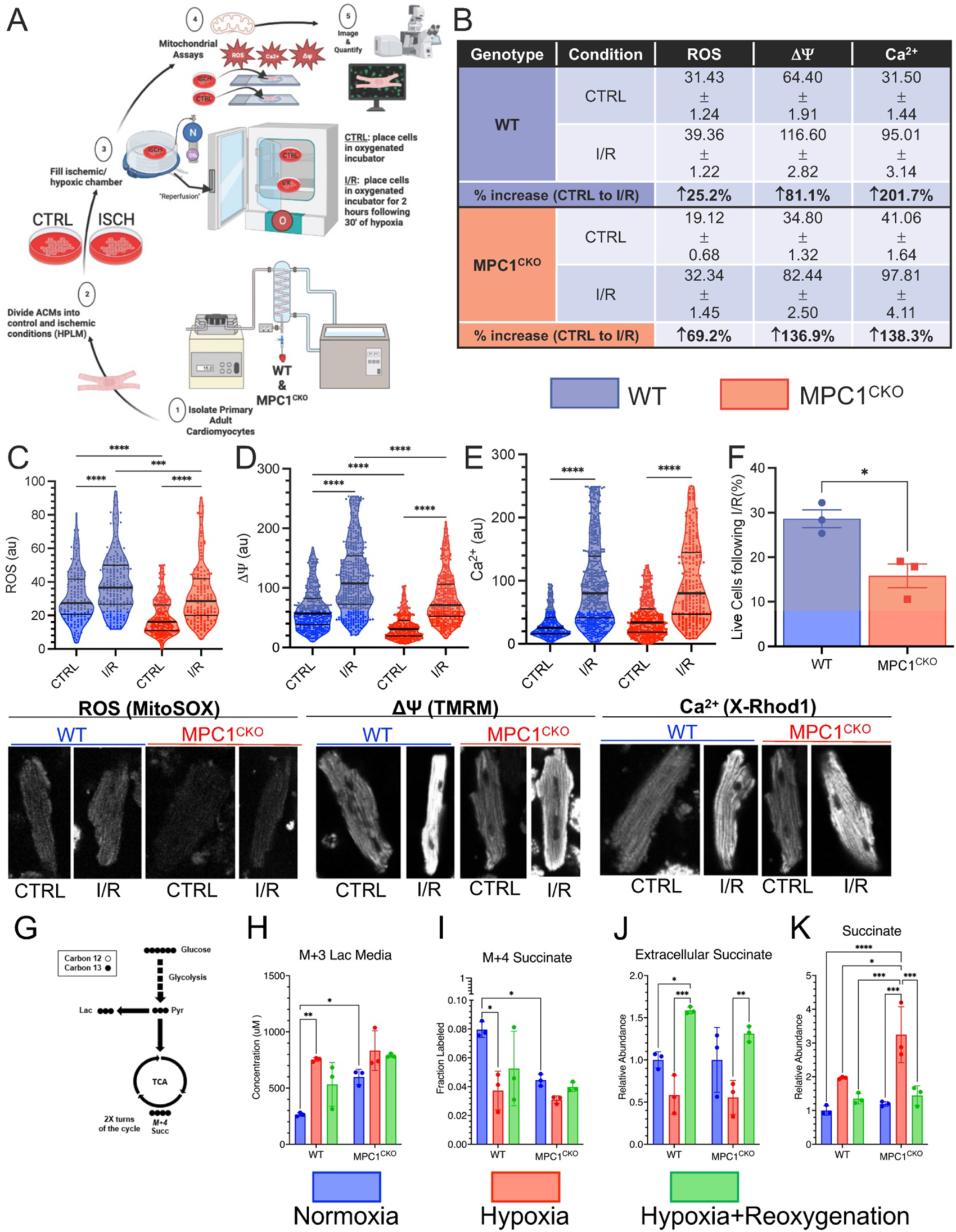
MPC1^CKO^ cardiomyocytes under simulated I/R exhibit greater mitochondrial damage, increased cell death, and more lactate/succinate flux. **A)** Schematic workflow for in vitro adult cardiomyocyte ischemic-reperfusion (hypoxia-reoxygenation) injury. **B)** Percent change in ROS, Δψ, and Ca^2+^ between WT and MPC1^CKO^ from CTRL to I/R conditions following in-vitro I/R. **C-E)** Mitochondrial ROS, membrane potential; Δψ, Ca^2+^ following I/R showing increases from control cells (CTRL: baseline) to I/R, with greater changes in the MPC1^CKO^ cells. **F)** Percentage of live cells following I/R injury. **G)** Diagram of [U-^13^C]glucose tracing in primary adult cardiomyocytes. Media M+3 lactate from shows MPC1^CKO^ cells efflux more lactate from glucose into the media at baseline. **I)** Cellular M+4 succinate labeling from [U-^13^C]glucose treated cells **J)** Extracellular succinate measured during reperfusion. **K)** Relative intracellular abundance of succinate during ischemia in WT and MPC1^CKO^. Unpaired t-tests and 2-way ANOVAs were used for statistical analysis between WT (n=3) and MPC1^CKO^ (n=3), and ischemic and non-ischemic tissue. * = p<0.05, ** = p<0.01, *** = p<0.001, **** = p<0.0001, ns=non-significant.

To understand how changes in metabolism were associated with the mitochondrial phenotypes we observed, we performed uniform [U-^13^C]glucose tracing in ACMs (Figure 3G). ACMs were exposed to hypoxia only, hypoxia + reoxygenation, or no treatment (control). In WT ACMs, as expected, hypoxia induced a large increase in glycolytic flux as assessed by media M+3 lactate excretion and this was partially reversed upon re-oxygenation (Figure 3H). In contrast, in normoxia, MPC1^CKO^ cardiomyocytes efflux more M+3 lactate into the media when compared to WT ACMs and this was not significantly increased by hypoxia. Hypoxia suppressed M+4 succinate production in WT ACMs, but this was already depressed in MPC1^CKO^ ACMs, and not further reduced by hypoxia (Figure 3I). Succinate is known to accumulate acutely in hypoxic cardiomyocytes and is associated with ROS injury upon reperfusion. Extracellular succinate accumulation was similar between genotypes in normoxia and decreased during hypoxia. Upon re-oxygenation, succinate was exported into the extracellular space similarly in both groups (Figure 3J). However, following hypoxia, the MPC1^CKO^ cardiomyocytes accumulated significantly higher levels of intracellular succinate compared to WT cells which returns to near baseline levels in both groups after 2 hours of reoxygenation (Figure 3K). In summary, lactate secretion is increased and glucose oxidation decreased in MPC1^CKO^ cells, and these parameters are no longer responsive to changes in oxygen tension. Total intracellular succinate accumulates to a higher level in MPC1^CKO^ cells, raising the possibility that these cells are subject to greater metabolic reperfusion injury.

### Pharmacological inhibition of MCT4 attenuates I/R injury

The upregulation of MCT4 accompanied by a simultaneous reduction of glucose oxidation and increase in lactate efflux in WT hearts and ACMs subjected to I/R led us to investigate whether this metabolic adaptation was actually deleterious. We hypothesized that redirecting glucose carbon flux away from lactate secretion and towards oxidation would improve mitochondrial substrate availability and mitigate reperfusion damage. To test this, we employed a selective MCT4 inhibitor (VB124) at the time of reperfusion to block MCT4-mediated lactate export. By increasing intracellular lactate levels, the lactate dehydrogenase (LDH) equilibrium is altered, and through mass-action pyruvate is shunted via the MPC towards mitochondrial oxidation. We previously demonstrated that MCT4 inhibition with VB124 can prevent cardiomyocyte hypertrophy ^18^.

Using our in vitro simulated I/R system, we treated ACMs from WT mice with VB124 at the time of reoxygenation. At baseline, there was minimal ROS emission and following hypoxia+reoxygenation ROS increased. However, administration of VB124 at the time of reoxygenation blunted ROS levels following hypoxia+reoxygenation (Figure 4A). We witnessed a similar pattern on membrane potential, with low Δψ at baseline and an elevation following hypoxia+reoxygenation, but this increase was also mitigated by VB124 (Figure 4B). We observed the same pattern for Ca^2+^ (Figure 4C). We then measured the percentage of live cells (Figure 4D) in normal and hypoxia+reoxygenation injury conditions and observed enhanced cell death after hypoxia+reoxygenation that was rescued by VB124 administration. Next, we examined if MCT4 inhibition successfully increased pyruvate oxidation. Following hypoxia+reoxygenation, ACMs treated with VB124 and [U-^13^C]glucose, showed a significantly higher fractional labeling of M+2 citrate when compared to vehicle-treated control cells (Figure 4E) indicating an increased contribution from glucose carbon.

**Figure 4:**
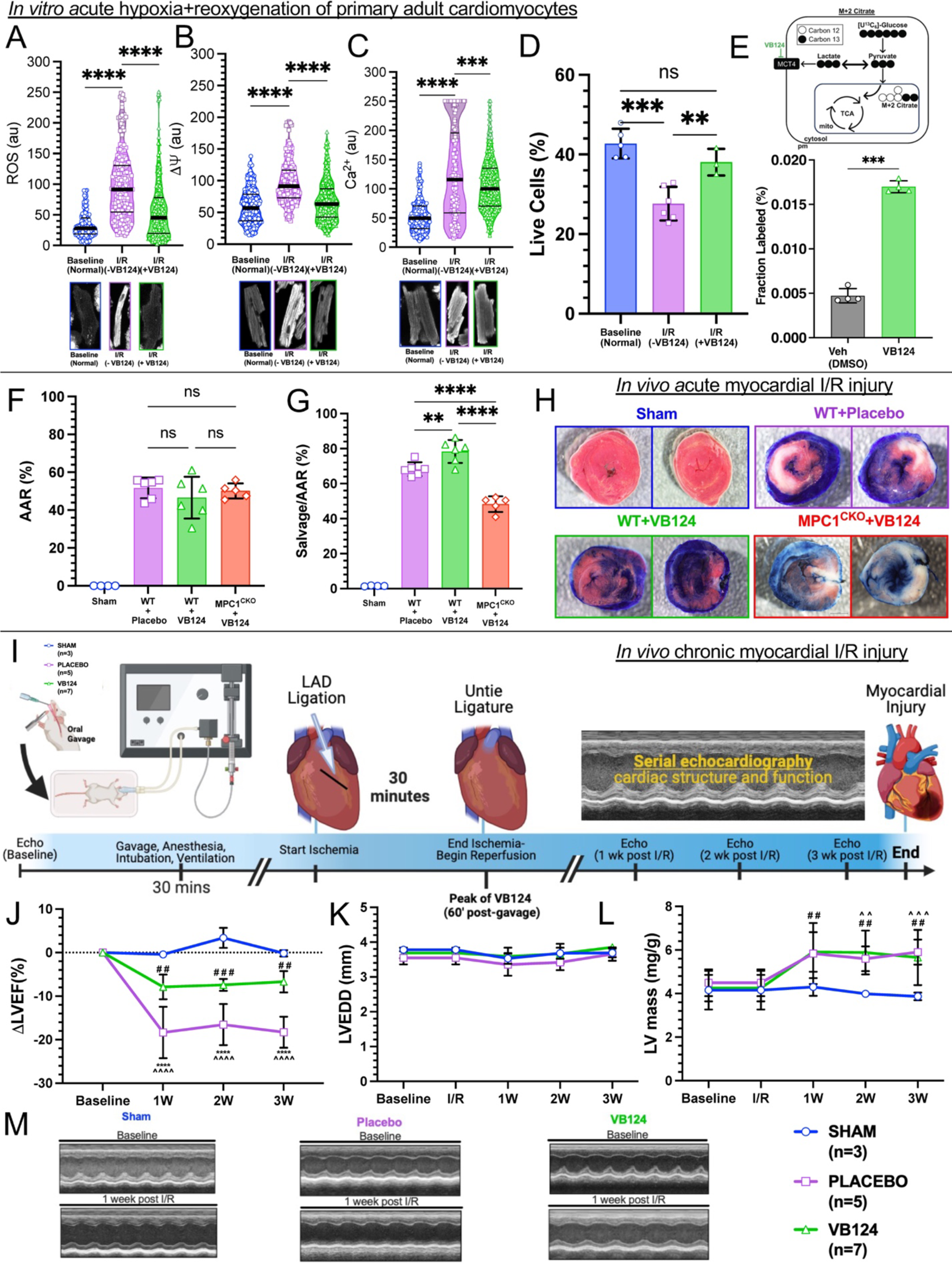
*In vitro* and *in vivo* MCT4 inhibition alleviates I/R injury. **A)** Mitochondrial ROS **(B)**, membrane potential; Δψ **(C),** Ca^2+^ following hypoxia+reoxygenation and the administration of VB124 showing increases from baseline to I/R, and then improvement with VB124 (normal, blue: n=5, -VB124, purple: n=6, +VB124, green: n=3). **D)** VB124 increases the cell viability and mitigates cell death following I/R injury. **E)** M+2 citrate labeling from [U-^13^C]glucose tracing in primary adult cardiomyocytes following hypoxia+reoxygenation with MCT4-inhibition (VB124, green) or Vehicle (DMSO, grey) controls (n=4, each). **F)** Following I/R injury, the area at risk (AAR %) is non-significant, indicating similar initial ischemic injury between placebo, +VB124, and MPC1^CKO^+VB124 groups (Sham: n=4, Placebo: n=6, VB124: n=6, MPC1^CKO^+VB124: n=5). **G)** Within the area at risk, myocardial salvage is significantly increased in the C57BL/6J mice (n=6), but not the MPC1^CKO^ (n=4) gavaged with VB124, after I/R injury when compared to placebo (n=6). **H)** Representative images of myocardial salvage, and necrosis following I/R injury in sham (n=4), placebo (n=6), +VB124 (n=6), and MPC1^CKO^+VB124 (n=5) groups. **I)** Schematic representation of *in vivo* chronic myocardial I/R injury with the placebo (methylcellulose) or VB124 oral gavage. **J-M)** Following I/R injury, serial echocardiography was performed to assess cardiac function (left ventricular ejection fraction; LVEF) and structure (left ventricular end-diastolic diameter; LVEDD and left ventricular mass; LV mass). Unpaired t-tests and 1-way ANOVAs were used for statistical analysis. Panels A-G) ** = p<0.01, *** = p<0.001, **** = p<0.0001, ns=non-significant. Panel J) **** = p<0.0001, Placebo compared to VB124. Panel L) ## = p<0.01, VB124 compared to Sham. ∧∧ = p<0.01, Placebo compared to Sham. ∧∧∧ = p<0.001, Placebo compared to Sham. Values are represented as mean±SD.

Encouraged by our *in vitro* results, we returned to an *in vivo* model of I/R injury and administered VB124 or placebo to healthy WT and MPC1^CKO^ mice (Supplemental Figure 4) via oral gavage. To determine the optimal timing for dosing, we orally gavaged mice with 30 mg/kg of VB124 and measured peak concentrations in the serum. VB124 reached a peak in the serum (Supplemental Figure 5, left) at 1-hour post-gavage and remained elevated for 4 hours. Therefore, we timed myocardial reperfusion for 1-hour post gavage in sham, placebo, WT+VB124, and MPC1^CKO^+VB124 mice. We collected serum and measured the concentration of VB124 by liquid chromatography-mass spectrometry (LC-MS) at baseline, the end of ischemia, and at the end of reperfusion. This analysis showed that the serum concentration of VB124 is highest in the WT and MPC1^CKO^ mice prior to reperfusion of the myocardium, and non-detectable in the sham, and placebo groups (Supplemental Figure 5, right). Controlling for infarct size, the AAR was similar between placebo, WT+VB124, and MPC1^CKO^+VB124 mice (Figure 4F). The myocardial salvage was then normalized to the AAR, and WT mice gavaged with VB124 experienced a statistically significant 15% increase in salvageable myocardium when compared to the placebo group, however this cardioprotective response was not seen in the MPC1^CKO^+VB124 mice (Figure 4G-H).

Finally, we tested whether inhibiting MCT4 could enhance long-term recovery from I/R. To test this, we replicated our I/R protocol administering either a single dose of placebo or VB124 (30mg/kg) via oral gavage prior to performing I/R. Unlike in the acute model, the mice were permitted to recover post-surgery, and we performed serial echocardiography over a three-week period to assess cardiac structure and function (Figure 4I). Initial assessments at baseline showed no significant differences between groups. Following 1, 2, and 3 weeks post-I/R injury, left ventricular ejection fraction (LVEF) was significantly reduced in the placebo and VB124 I/R groups compared to sham-operated mice, but treatment with VB124 mitigated this decrease (Figure 4J). There were no significant changes to left ventricular end-diastolic diameter (LVEDD) between groups (Figure 4K). LV-mass increased similarly after injury in both the placebo and VB124 groups, and at 3 weeks post-I/R both groups had significantly larger LV-mass compared to Sham (Figure 4L). Sham-operated mice remained free of any signs of cardiac injury. Our results show that acute MCT4 inhibition therapy suppresses increased ROS, Δψ, and Ca^2+^ following I/R injury, increases pyruvate flux to TCA cycle, and improves myocardial salvage leading to a sustained improvement in cardiac function.

## DISCUSSION

Despite decades of research, no effective therapies are available to treat myocardial I/R injury ^2^. Prior proposed interventions targeting metabolic aspects of I/R, such as protein kinase C inhibitors ^30^, and ATP-sensitive potassium channel openers ^33^, have not improved outcomes in the clinic, illustrating the need for new conceptual approaches. The MPC complex is responsible for transporting pyruvate, the end product of glycolysis, into the mitochondria for oxidation coupled to the synthesis of ATP. The MPC is upregulated at both the transcript and protein level in the surviving myocardium post I/R injury, suggesting a new mechanism of cardioprotection via enhanced pyruvate oxidation ^22^. Similarly, it was recently reported that the downregulation of *MPC2* in the brain aggravates neuronal injury following cerebral I/R ^34^. Our group and others recently discovered that mice subjected to inducible heart-specific deletion of *MPC1* spontaneously develop hypertrophic cardiomyopathy, become moribund, and perish at 16 weeks post induction due to HF symptoms ^18, 23–25^. Based on the in vivo data showing a cardioprotective role of the MPC, we hypothesized that MPC1^CKO^ hearts would suffer from a worse I/R injury when exposed to a cardiac insult prior to the development of HF symptoms.

### MPC1^cko^ increases I/R injury

We subjected MPC1^CKO^ mice 4 weeks post-injection with tamoxifen to acute I/R injury. Knockout animals suffered worse infarcts with 20% less myocardial salvage and more necrosis compared to littermate controls (Figure 1). Within the myocardium of MPC1^CKO^ animals, we observed an upregulation of several deleterious pathways and transcripts in the MPC1^CKO^ ischemic myocardium after I/R injury (Figure 2E). We also observed increased expression of *Slc16a3* (MCT4) (Figure 2F-G). MCT4 facilitates lactate efflux from cardiomyocytes, and its increased expression suggests that, in the absence of the MPC, pyruvate is being converted to lactate within the cytoplasm of these cells and exported. Of note, we also observed an injurious transcriptional phenotype within the non-ischemic myocardium of the MPC1^CKO^ hearts, which indicate that the entire heart may undergo transcriptional changes following acute I/R injury.

Turning to an *in vitro* model to better characterize the cellular mechanisms underlying the genotype-specific responses to I/R injury, we exposed isolated ACMs to hypoxia before restoring normoxia and quantified ROS, Δψ and Ca^2+^ as well as glucose metabolism using stable isotope tracers. MPC1^CKO^ ACMs had reduced baselines levels of ROS and Δψ compared to WT suggesting that MPC1^CKO^ mitochondria may already exhibit an impaired energetic state. We can’t rule out that this baseline difference was exacerbated by the media conditions ACMs are cultured in, as these mice show no cardiac phenotype at this timepoint. The decompensated state in the MPC1^CKO^ mitochondria proved to be pathological as they demonstrated significantly more cell death with greater changes from baseline to stressed conditions when analyzing ROS and Δψ, but not Ca^2+^ (Figure 3). [U-^13^C]glucose labeling experiments demonstrated that, even in normoxia, MPC1^CKO^ have increased lactate excretion. During hypoxia the MPC1^CKO^ ACMs accumulated higher levels of succinate compared to WT cells, indicating impaired succinate oxidation/excretion following I/R injury, which has been shown to be detrimental^12^. Our findings that absent pyruvate oxidation, cells show impaired mitochondrial responses to I/R injury strongly implicate pyruvate as a critical metabolite in conditions of hypoxia.

### MCT4 inhibition can alleviate I/R injury

Pyruvate exists in an equilibrium with lactate catalyzed by the enzyme lactate dehydrogenase. Lactate efflux from the cell is accomplished by the primary lactate exporter MCT4. Although MCT4 typically exhibits low expression in cardiomyocytes, it may be upregulated during pathological/stressful conditions that increase glucose consumption to fuel glycolytic metabolism in oxygen limiting conditions such as hypoxia, or extreme exercise ^35, 36^. While increased MCT4 may be an acutely adaptive response to heart injury, enhancing lactate efflux may actually be counterproductive as it facilitates increased LDH reaction flux thereby siphoning pyruvate away from oxidation and enabling higher rates of glycolytic flux through rapid regeneration of cytoplasmic NAD+. In support of this idea, MCT4 inhibition therapy in mouse models has been established to prevent cardiomyocyte hypertrophy, and attenuate cardiac injury following pulmonary embolism and chronic pressure overload ^18, 37, 38^.

Upon I/R, we observed enhanced expression of MCT4 in MPC1^CKO^ injured hearts compared to WT, likely due to high levels of intracellular lactate production. Returning to our in vitro hypoxia+reoxygenation model, administration of the MCT4 inhibitor VB124 at the time of reoxygenation mitigated the harmful effects of I/R injury, increased pyruvate oxidation and rescued cardiomyocyte death (Figure 4A-E). We translated these findings in vivo in our acute I/R mouse model. We showed that VB124 treatment during I/R resulted in a 15% increase in myocardial salvage following injury when compared to placebo-treated mice (Figure 4F-H). Interestingly, MPC1^CKO^ mice gavaged with VB124 did not show improvements to myocardial salvage following I/R injury, suggesting that MPC-mediated pyruvate transport is necessary for the cardiac benefit of MCT4 inhibition. Finally, we sought to understand whether these improvements in acute markers of heart injury would lead to long-term increases in cardiac function. Single-dose treatment with VB124 at the time of injury conferred durable cardioprotection, as shown by the preservation of cardiac function (LVEF) when compared to placebo-treated mice at 3 weeks post injury (Figure 4J). Interestingly, mice treated with either placebo or VB124 showed similar increases in normalized LV mass over body weight (mg/g) indicating that the hypertrophic response was not attenuated by MCT4 inhibition, but only the functional consequences, perhaps reflecting a reduced initial injury (Figure 4L).

These results show that MCT4 inhibition can be cardioprotective following I/R injury. Our data suggest that inhibiting MCT4 has the potential to alleviate acute and chronic I/R injury by redirecting the fate of pyruvate and improving mitochondrial oxidative metabolism. We hypothesize this may in part be due to the ability for MCT4 inhibition to increase mitochondrial pyruvate metabolism toward ATP production and away from other potentially deleterious metabolic pathways. Moreover, pyruvate has been shown to neutralize ROS, by scavenging free radical species ^39–41^. This additional mechanism may protect the myocardium from oxidative stress, and support the transition towards healthy cardiac metabolism following injury, however more research is needed to support this hypothesis.

### Conclusions and Perspectives

Despite decades of research, there are currently no effective clinical therapies in use today that alleviate I/R injury to the heart. Here, we present data to support pyruvate oxidation and the MPC to be cardioprotective in the setting of myocardial I/R injury. Our data in multiple systems show that the use of an MCT4 inhibitor could improve outcomes after I/R injury. Therefore, we suggest that interventions that promote pyruvate oxidation could alleviate I/R injury. Future research should focus on the exact mechanisms of how MCT4 inhibition can salvage the myocardium and address the efficacy, dose-response, and therapeutic window of this potential.

### Limitations

#### In vitro

Simulated ischemia-reperfusion systems with hypoxia and reoxygenation are commonly used to study the mechanisms associated with I/R injury. However, the in vitro system has limitations that need to be taken into consideration. Chiefly, in vitro ACM systems lack mechanical stress and the complexity of diverse cell types which may influence results. Moreover, the variability in the quality and health of primary adult cardiomyocyte during isolation and culture can introduce variation in results.

#### In vivo

The left anterior descending (LAD) artery occlusion is a widely used method to study I/R injury. In this study, the invasiveness of the in vivo I/R procedure requires surgery and can be associated with a risk of complications such as bleeding and mortality. For our tissue staining we were able to control for infarct size, however, the size of the infarct can vary depending upon the location of the occlusion, and the extent of collateral branches that may be present. Additionally, the 30-minute duration of ischemia that we used may not accurately translate to longer periods of ischemia that may occur in patients.

## KEY RESOURCES TABLE

| Reagent or resource | Source | Identifier |
| --- | --- | --- |
| **Antibodies** |  |  |
| MPC1 | Cell Signaling Technology | 14462S |
| MCT4 | Santa Cruz Biotechnology | Sc-376140 |
| **Biological Samples** |  |  |
| **Chemicals, Peptides, and Recombinant Proteins** |  |  |
| 0.9 % NaCl TC grade | Sigma Aldrich | S8776-100ML |
| VB124 | Vettore Biosciences | N/A |
| LC/MS grade water | Fisher Scientific | W6-1 |
| LC/MS grade methanol | Fisher Scientific | A456-1 |
| [U-<sup>13</sup>C]glucose | Cambridge | CLM-1396 |
| Fetal Bovine Serum, dialyzed,<br>One Shot™ | Thermo Scientific | A3382001 |
| Primocin | InvivoGen | Ant-pm-2 |
| Natural Mouse Laminin | Thermo Scientific | 23017-015 |
| HEPES | Thermo Scientific | 15630080 |
| Insulin-Transferrin-Selenium | Thermo Scientific | 41400045 |
| 2,3-Butanedione monoxime | Sigma Aldrich | B0753-25G |
| Bovine Serum Albumin | Fisher Scientific | BP9705100 |
| **Recombinant DNA** |  |  |
| **Critical Commercial Assays** |  |  |
| Pierce BCA protein assay kit | Thermo Scientific | 23225 |
| miRNeasyMini Kit | Qiagen | 74104 |
| RNAase free DNAase kit | Qiagen | 79254 |
| TruSeq Stranded RNA kit with<br>Ribo-Zero Gold | Illumina | RS-122-2301 |
| QIAamp® DNA Blood Maxi kit | Qiagen | 51192 |
| **Experimental models: Cell Lines** |  |  |
| **Experimental Models: Organisms/Strains** |  |  |
| Mouse: C57BL/6J | Jackson Laboratories | Cat# 000664 |
| Mouse: *MPC1<sup>CKO</sup>* | This work | N/A |
| **Software and Algorithms** |  |  |
| Prism 10 | GraphPad Software | https://graphpad.com |
| LI-COR Image Studio | LI-COR Biosciences | https://licor.com |
| **Other** |  |  |
| Waters GCT Premier Mass spectrometer | Waters |  |
| Agilent 6890 gas chromatograph | Agilent |  |
| Gerstel MPS2 autosampler | Gerstel |  |
| HiSeq 2500 | Illumina |  |

## RESOURCE AVAILABILITY

### Lead Contact

Further information and requests for resources and reagents should be directed and will be fulfilled by the Lead Contacts, Stavros G. Drakos, Jared Rutter, and Gregory Ducker

### Materials availability

Mouse and other materials generated in this study are available for any researcher upon request.

### Data availability

Sequencing data is available upon request.

## EXPERIMENTAL MODEL AND DETAILS

### Animals and animal care

All animal experiments conducted in this study were approved by and performed in accordance with the Institutional Animal Care and Use Committee (IACUC) at the University of Utah. To generate adult cardiac-specific MPC1-knock out mice, a C57BL/6 MPC1 fl/fl mice were bred to αMHC-MerCreMer mice. The resulting MPC1 fl/+; αMHC-MerCreMer were then bred to each other, resulting in the production of MPC1 fl/fl; αMHC-MerCreMer as well as their WT littermates, which were used as controls. To achieve adult cardiac-specific MPC1 knockout and WT littermates, 8-week-old MPC1 fl/fl; αMHC-MerCreMer and their WT littermates were intraperitoneally injected with 40 mg/kg of tamoxifen for three consecutive days.

### In-vivo ischemic-reperfusion injury

The experimental protocol was performed using 12–14-week-old mice (4 weeks post tamoxifen injection) prior to the development of HF symptoms, weighing 20-30 g and their WT littermate controls. At least 2 days prior to the scheduled surgery date, special attention was given to guarantee the cleanliness and the maximum sterility of the operation area and the well-being of mice. Consequently, clean surgical gown, and sterile gloves were worn. The cleanness of the surgery area was checked and assured by bleaching the sterile area utilized as surgical zone and by using sterile drapes to delimit it. Finally, all surgery instruments were autoclaved and sterilized.

On the surgery day, mice were anesthetized with 2-3% isoflurane inhalation, orally intubated with a 22 GIV catheter, and artificially ventilated with a small rodent respirator (Harvard Instrument). The level of anesthesia was monitored by testing for corneal reflex and muscle tone. The animals were ventilated at a rate of 100/minute with tidal volume adjusted to produce peak pressures of 15-20 mmHg. Anesthetized mice were placed on a circulating warming pad (38° C) in the supine position. Chest hairs were removed with a topical depilatory agent. Depilatory cream was then rinsed thoroughly with water or saline prior to surgical scrub to prevent skin irritation. The skin was then prepared with betadine and alcohol. A thoracotomy was performed to visualize the anterior surface of the heart. Next, a suitable retractor was used to retract the thoracic wall to improve visualization and accessibility. The pericardium was removed with the use of forceps and scissors. Under the dissecting microscope the left anterior descending coronary artery (LAD) was visualized, and a 6-0 silk suture and the needle were inserted under the artery, as the needle enters the myocardium, we were careful to avoid bleeding. A loose double knot was then made with the suture, leaving a 2-3 mm diameter loop through, then a 2-3 mm long piece of PE-10 tubing was placed. The tubing was soaked in 100% ethanol for 24 hours and rinsed free of alcohol with sterile water before use. The loop around the artery and tubing was tightened for 30 min to ensure coronary artery occlusion. The characteristic paler color appearance of the anterior wall of the LV appeared within a few seconds following LAD ligation which we used as confirmation that the vessel was occluded. After the 30 minutes of ischemia is complete the knot was untied and the PE-10 tubing removed. The reperfusion was confirmed by observing a return of red color to the anterior wall of the LV after a few seconds. Then, the reperfusion continued for 120 mins (2 hours). After completion of the ischemia reperfusion (I/R) injury, the coronary artery was briefly re-occluded, and Evans blue was injected into the left atrium to demarcate the area at risk (AAR). Hearts were then excised, perfused with 0.9% saline, weighed, and then cut into 1-mm-thick transverse slices using a vibratome device, or separated into ischemic and non-ischemic tissue samples to be flash frozen in liquid nitrogen. The slices were then incubated in 1% triphenyltetrazolium chloride (TTC) in sodium phosphate buffer (pH 7.4) at 38°C for 20 min. TTC stains the non-infarcted myocardium. The slices were then immersed in 10% formalin to enhance the contrast between stained (viable) and unstained (necrotic) tissue. Tissue sections were imaged the following day and myocardial salvaged was quantified by independent researchers who were blinded to the treatment groups using ImageJ (NIH Software).

### Echocardiography

Under 1-3% isoflurane anesthesia (VetOne Fluriso, NDC 13985-528-60), mice underwent echocardiography using the Vevo imaging system. Serial echocardiograms included 2D long-axis and short-axis views for analysis with Vevo strain software (version 3.1.1). Analysis was based on two consecutive cardiac cycles were used for all the measurements. Limb leads were used to record an electrocardiogram (ECG). To ensure objectivity, all echocardiography data were analyzed an independent researcher who was blinded to the treatment groups.

### Primary adult cardiomyocyte isolation and culture

Adult mice were anesthetized and the heart was excised, attached to an aortic cannula, and perfused with solutions held at 37 degrees C, pH 7.3. Perfusion with a 0 mM Ca^2+^ solution for 5 minutes was followed by 15 minutes of perfusion with the same solution containing 1 mg/mL collagenase and 0.1 mg/mL protease. The heart was then perfused for 1 minute with a “stopping solution” (the same solution containing 20% serum and 0.2 mM CaCl_2_). All perfusions were performed at a flowrate of 2 mL/min. The atria were removed, and the ventricles minced and then filtered through a nylon mesh. Cells were stored at 37°C in culture media. All cardiomyocytes used in this study were rod-shaped, had well defined striations, and did not contract spontaneously.

Following isolation, primary adult cardiomyocytes were enriched by gravity sedimentation in increasing concentrations of Ca^2+^ plating media. Cells were plated on Poly-L-lysine coated Petri dishes an allowed to adhere for at least 1 hour in the incubator. Cells were then cultured in a final Ca^2+^ concentration of 1mM.

### In-vitro ischemic-reperfusion injury (hypoxia/reoxygenation)

Primary adult cardiomyocytes were isolated using the method described above and then placed in an airtight container with a constant gas flow of 95% nitrogen, and 5% carbon dioxide at 37°C. This process ensured the complete removal of oxygen from within the sealed ischemia chamber. For ROS, Δψ, and Ca^2+^ measurements the ischemia lasted for 2 hours. Following ischemia, cells were removed from the sealed container and placed in the oxygenated incubator alongside the control cells, and measurements were taken following 2 hours of reperfusion. Control cells were kept at 37°C for the same amount of time that the experimental cells were exposed to the I/R conditions.

## METHOD DETAILS

### Tissue lysates

Proteins from mouse heart were extracted from either ischemic, or non-ischemic tissue samples. Briefly, tissue was placed in a 2mL Eppendorf tube with 1X RIPA lysis buffer, which contained HALT protease and phosphatase inhibitor cocktail solution (Pierce, Rockford, IL). A bead-homogenizer (Tissue Lyser II, Qiagen) was used to homogenize the tissue samples and the lysates were incubated on ice for one hour. Samples were then centrifuged at 14,000 g for 30 minutes at 4°C. Next, the supernatant was collected, and the Pierce BCA Protein Assay Kit (Thermo Scientific, Rockford, IL) was used for protein estimation, before equal volumes of 2x sample buffer containing 1 mM DTT was added to the samples.

### Western blotting

Following estimation of protein concentration, samples were boiled for 5 minutes at 95°C. 30-50 μg of total protein lysate was resolved on SDS polyacrylamide gel according to the standard procedure of 20 mA per gel and blotted onto a nitrocellulose membrane 0.45 μm (GE Healthcare) via Mini Trans-blot module (Bio-Rad) at a constant voltage of 100 V for 2 hours. Next, membranes were blocked with 5% proteomic grade non-fat dried milk (NFDM) for 1 hour and then the membranes were incubated overnight in 5% bovine serum albumin. Antibodies probing for MPC1 (cell signaling 1:1000), and MCT4 (Santa Cruz 1:1000) with a loading control of either Vinculin (cell signaling 1:1000) or HSP60 (cell signaling 1:1000) were used. Following incubation and on the next day, membranes were washed with TBS-T and placed in an incubation with a fluorophore conjugated secondary antibody (Rockland Immunochemical, 1:10,000) in 1% NFDM with TBS-T for 1 hour. Next, membranes were washed again with TBS-T and fluorescence was detected using an Odyssey CLx imaging system (LI-COR Biosciences).

### Gene expression analysis

Total RNA from mouse hearts (normal, ischemic, or non-ischemic) was isolated using RNeasy Mini kits (Qiagen), according to the manufacturer’s instructions. Next, cDNA was synthesized using a cDNA Reverse Transcriptase Kit (New England Biolabs). TaqMan-based real time quantitative polymerase chain reactions (qRT-PCR) were then performed using a QuantStudio 7 Pro Real-Time PCR System (ThermoFisher). The housekeeping gene *Vinculin* was used as an internal control for cDNA quantification and normalization of gene amplified products.

### Mouse RNA-sequencing

Bulk RNA was isolated from mouse hearts (normal, ischemic, or non-ischemic) using the miRNeasy Mini Kit (Qiagen). Total RNA samples (100-500 ng) were hybridized with Ribo-Zero Gold (Illumina) to deplete cytoplasmic and mitochondrial rRNA from the samples. Stranded RNA sequencing libraries were prepared as described using the Illumina TruSeq stranded Total RNA Library Prep Gold kit (20020598) with TruSeq RNA UD Indexes (20022371). Purified libraries were qualified on an Agilent Technologies 2200 TapeStation using a D1000 ScreenTape assay (cat# 5067-5582 and 5067-5583). The molarity of adaptor-modified molecules was defined by qRT-PCR using the Kapa Biosystems Kapa Library Quant Kit (cat# KK4824). Individual libraries were normalized to 1.30 nM in preparation for Illumina sequence analysis. Sequencing libraries (1.3 nM) were chemically denatured and applied to an Illumina NovaSeq flow cell using the NovaSeq XP chemistry workflow (20021664). Following transfer of the flowcell to an Illumina NovaSeq instrument, a 2 x 51 cycle paired end sequence run was performed using a NovaSeq S1 reagent kit (20027465).

### RNA-seq analysis and bioinformatics

RNA-seq analysis was conducted with the High-Throughput Genomics and Bioinformatics Analysis Shared Resource at Huntsman Cancer Institute at the University of Utah. Briefly, the mouse GRCm38 FASTA and GTF files were downloaded from Ensembl release 96 and the reference database was created using STAR version 2.7.0f with splice junctions optimized for 50 base-pair reads. Optical duplicates were removed from the paired end FASTQ files using BBMap’s Clumpify utility (v38.34) and reads were trimmed of adaptors using cutadapt 1.16. The trimmed reads were aligned to the reference database using STAR in two-pass mode to output a BAM file sorted by coordinates. Mapped reads were assigned to annotated genes in the GTF file using featureCounts version 1.6.3. The output files from cutadapt, FastQC, Picard CollectRnaSeqMetrics, STAR, and featureCounts were summarized using MultiQC to check for any sample outliers. Differentially expressed genes with at least 95 read counts across all samples for a given tissue (normal, ischemic, or non-ischemic) were identified using DESeq2 version 1.24.0. Genes were then filtered using the criteria FDR < 0.01, absolute log2 fold change > 0.0.585, (fold change > 1.516). Up- and downregulated gene sets from each tissue were then analyzed for enriched GO terms using Gprofiler with a Benjamini-Hochberg FDR p-value correction thresholded at < 0.01. RNA-seq read counts were RPKM normalized for heatmap generation. Heat-maps were generated by gene-standardizing the RPKM gene values (mean = 0, stdev = 1) and plotting using XPRESSplot v0.2.2, Matplotlib, and Seaborn.

### Metabolite extraction

The protocols for metabolite extraction from cultured cells have been described previously. However, following the *in vitro* ischemic-reperfusion injury (hypoxia/reoxygenation) the media was collected and aspirated. Adherent cells were washed with cold 0.9% sterile saline which was chilled on ice. Next, 3 mL of extraction solvent (80% methanol/water) cooled to -80°C was added to the dish and then the dish was transferred to a -80°C freezer. Cells were then scraped into the extraction solvent on dry ice. All metabolite extracts were centrifuged at 20,000 rcf at 4°C for 10 minutes. Each sample was then transferred to a new 1.5 mL tube. Lastly, the solvent in each sample was evaporated in a speed vacuum overnight and stored at -80°C until they were analyzed via mass spec.

### [U-^13^C]Glucose labeling

Following cardiomyocyte isolation, the cells were placed in 10 cm plates with media where the normal glucose was replaced with ^13^C_6_-L-glucose (Cambridge Isotope Laboratories). Cells were allowed to incubate in the ^13^C_6_-L-glucose prior to the *in vitro* ischemic-reperfusion injury (hypoxia/reoxygenation) experiments. Following I/R injury, metabolites were extracted as described above and the data was corrected for naturally occurring ^13^C isotope abundance before analysis.

### Statistical Analysis

We have presented data with either the mean ± standard deviation (SD) or ± standard error of the mean (±SEM). For our statistical evaluations, we utilized GraphPad Prism software, version 10.0.0. We applied two-tailed tests for all comparative analyses. When assessing two distinct groups, we employed the unpaired Student’s t-test and the two-tailed Mann-Whitney test. For evaluating serial echocardiographic data and multiple group comparisons, we conducted multiple t-tests and ANOVAs, respectively.

## Supporting information

Supplement_Visker 2024

## Acknowledgements

We thank Chris Stubben for his data analysis and advice related to transcriptomics. The authors would also like to thank K. Mark Parnell with Vettore Biosciences for his expertise in working with VB124. We also thank members of the Drakos, Rutter, Ducker, and Chaudhuri labs for their assistance and helpful discussions. Additionally, we would like to thank the Nora Eccles Harrison Cardiovascular Research and Training Institute for their support of this research project. We acknowledge the following funding: the National Institutes of Health under Ruth L. Kirschstein National Research Service Award T32HL007576 from the National Heart, Lung, and Blood Institute of the National Institutes of Health to JRV, and Award Number K99HL168312 for A.A.C. J.R. is an investigator of the Howard Hughes Medical Institute. The content of this manuscript is sole the responsibility of the authors and does not necessarily represent the official views of the National Institute of Health (NIH).

## Author Contributions

Conceptualization, J.R.V., A.A.C, J.V., D.C., S.N., G.S.D, J.R., and S.G.D.; Methodology, Validation, Formal Analysis, and Investigation, J.R.V., A.A.C, J.V., D.E., D.C., T.S.S., R.H., J.L., H.K., and Y.H.,; Resources, D.C., D.P., G.S.D.; Data Curation, I.A., J.S.V., A.A.C., J.V., and D.E.; Writing – Original Draft, J.R.V., A.A.C., J.V., G.S.D., S.G.D, and J.R.; Writing – Review & Editing, J.R.V., A.A.C., J.V., G.S.D., J.R., S.G.D., and S.N.; Visualization, J.R.V., A.A.C., J.V., D.E., T.S.S., R.H., H.K., and Y.H..; Supervision, G.S.D., J.R., and S.G.D.; Funding Acquisition, J.R., and S.G.D.

