## Supplement_Visker 2024 for "Enhancing mitochondrial pyruvate metabolism ameliorates myocardial ischemic reperfusion injury"

### SUPPLEMENTAL MATERIALS

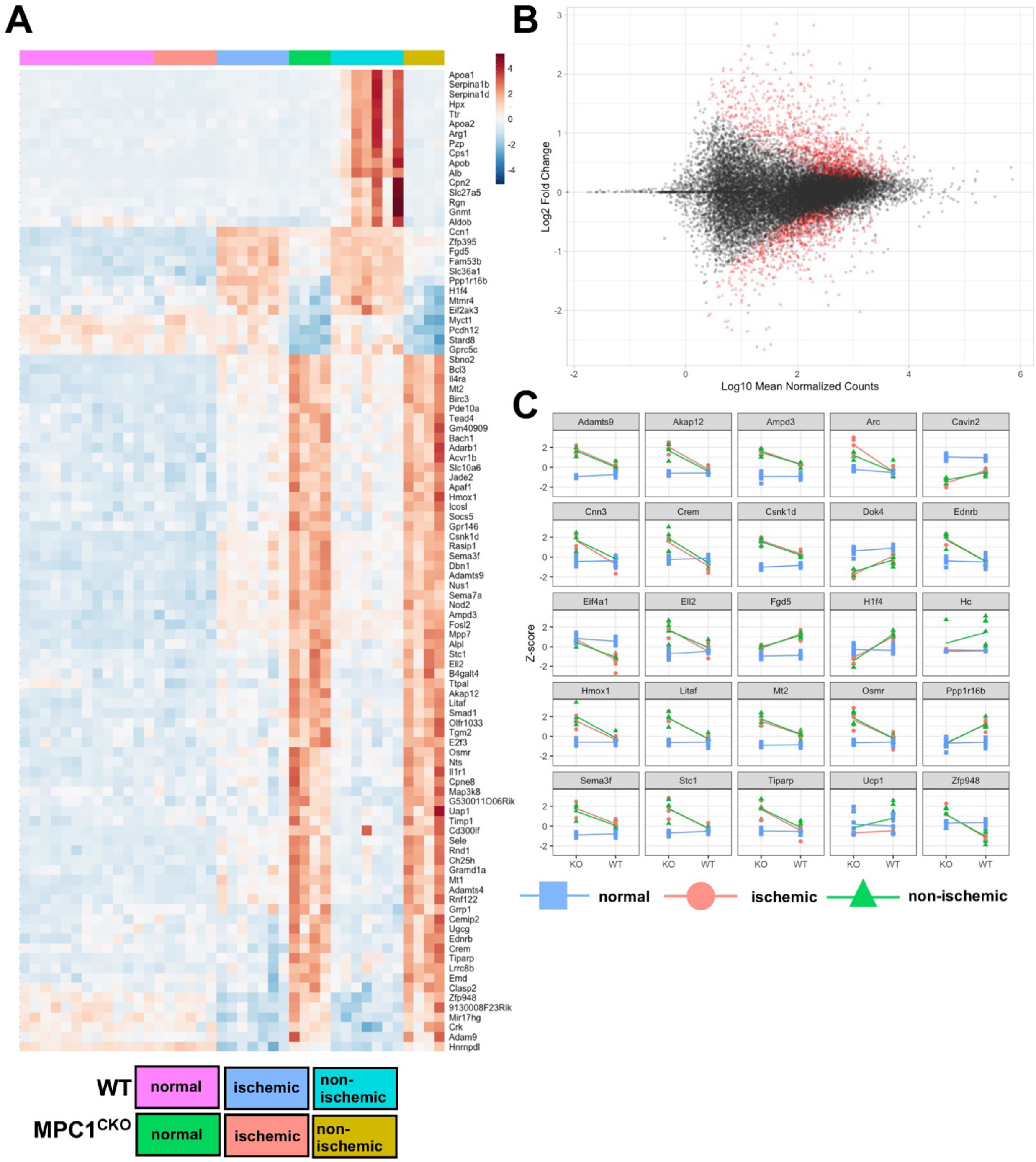

**Supplemental Figure 1: A)** Differential gene expression heatmap in I/R injury showing the global gene expression profiles of the top 100 differentially expressed genes in myocardial tissue from WT and $MPC1^{CKO}$ mice. Each row represents a gene, and each column corresponds to a tissue sample, categorized by genotype and myocardial tissue status: normal, ischemic, and non-ischemic. The color gradient, from blue to red, denotes the expression level from low to high (z-scores), showing gene expression patterns across different conditions and genotypes. The analysis was conducted using an adjusted p-value of less than 0.05 and a false discovery rate (FDR) of 5%. **B)** MA-plot showing the differential gene expression analysis between $MPC1^{CKO}$ and WT ischemic myocardium. The x-axis displays the Log10 mean normalized counts, which is the average expression of each gene across all samples, providing a measure of gene abundance. The y-axis shows the Log2 fold change, representing the change in gene expression between the two conditions. Points in red indicate the top 200 significant genes with differential expression, clustered and scaled by rows. **C)** Interaction plots in Ischemic, Non-Ischemic, and Normal Conditions across genotypes (WT or $MPC1^{CKO}$) showing the scaled relative log expression (rlog) values, represented as z-scores, for the top 25 statistically significant ($p<0.05$) genes comparing ischemic, non-ischemic, and normal conditions across WT and $MPC1^{CKO}$ mice.

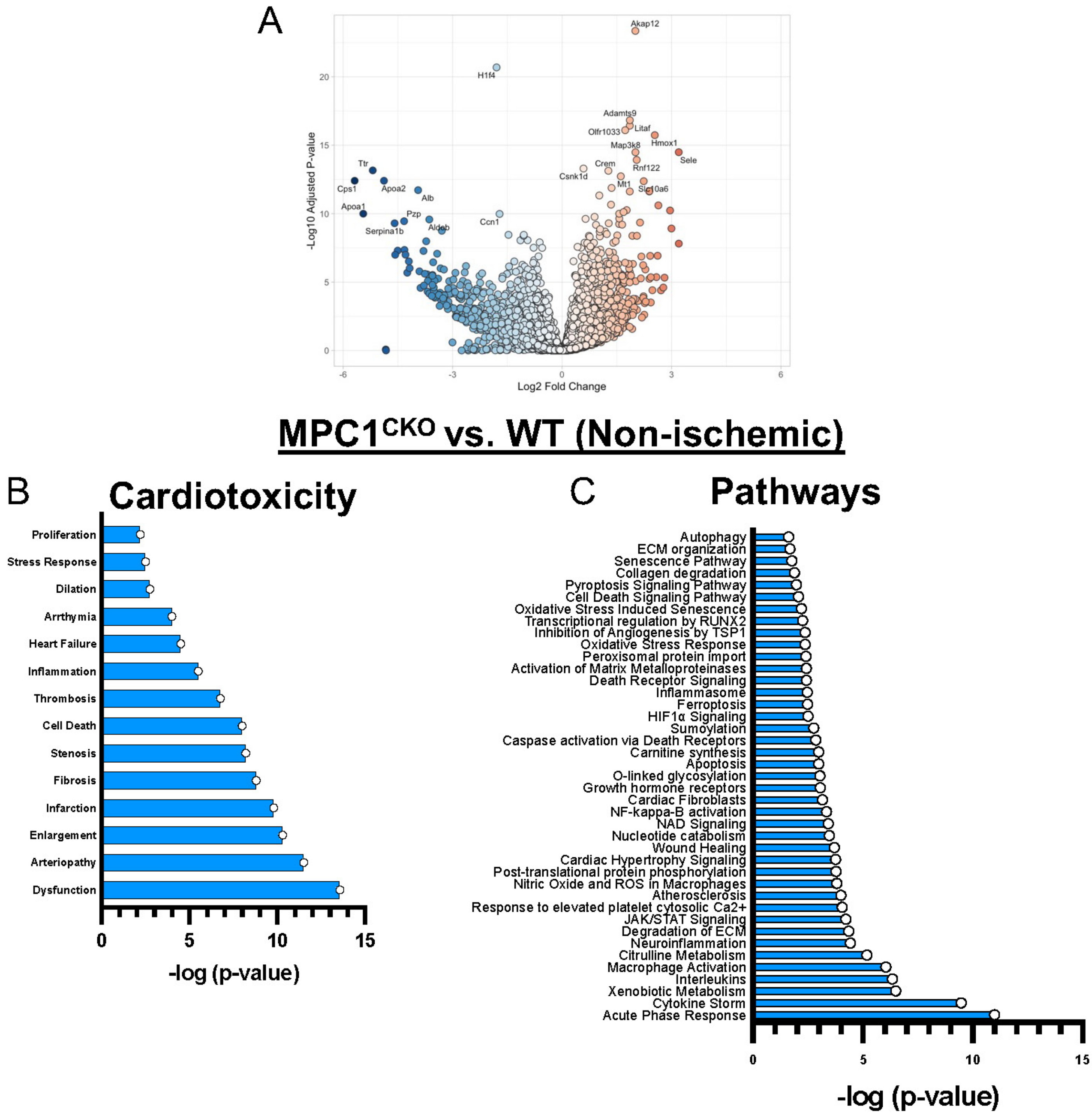
A
Akap12
H1f4
Adamts9
Olfr1033
Litaf
Hmox1
Map3k8
Sele
Crem
Rnf122
Csnk1d
Mt1
Slc10a6
Ttr
Cps1
Apoa2
Alb
Apoa1
Pzp
Aldob
Ccn1
Serpina1b
-Log10 Adjusted P-value
Log2 Fold Change
MPC1CKO vs. WT (Non-ischemic)
B
Cardiotoxicity
Proliferation
Stress Response
Dilation
Arrthymia
Heart Failure
Inflammation
Thrombosis
Cell Death
Stenosis
Fibrosis
Infarction
Enlargement
Arteriopathy
Dysfunction
-log (p-value)
C
Pathways
Autophagy
ECM organization
Senescence Pathway
Collagen degradation
Pyroptosis Signaling Pathway
Cell Death Signaling Pathway
Oxidative Stress Induced Senescence
Transcriptional regulation by RUNX2
Inhibition of Angiogenesis by TSP1
Oxidative Stress Response
Peroxisomal protein import
Activation of Matrix Metalloproteinases
Death Receptor Signaling
Inflammasome
Ferroptosis
HIF1α Signaling
Sumoylation
Caspase activation via Death Receptors
Carnitine synthesis
Apoptosis
O-linked glycosylation
Growth hormone receptors
Cardiac Fibroblasts
NF-kappa-B activation
NAD Signaling
Nucleotide catabolism
Wound Healing
Cardiac Hypertrophy Signaling
Post-translational protein phosphorylation
Nitric Oxide and ROS in Macrophages
Atherosclerosis
Response to elevated platelet cytosolic Ca2+
JAK/STAT Signaling
Degradation of ECM
Neuroinflammation
Citrulline Metabolism
Macrophage Activation
Interleukins
Xenobiotic Metabolism
Cytokine Storm
Acute Phase Response
-log (p-value)

**Supplemental Figure 2: A.)** Volcano plot showing differential expression of genes in non-ischemic heart tissue derived from WT and MPC1$^{CKO}$ heart tissue following I/R injury. Each dot corresponds to a specific gene. The x-axis shows the log2 fold change in gene expression, indicating the magnitude of upregulation (to the right: red) or downregulation (to the left: blue). The y-axis shows the negative logarithm (base 10) of the adjusted p-value, which represents the statistical significance of the gene expression change. The upregulated genes in the MPC1$^{CKO}$ non-ischemic tissue include markers such as Akap12, Adamts9, Hmox1, and Sele. Downregulated genes in the MPC1$^{CKO}$ non-ischemic tissue include Cps1, Apoc1, and Serpinab1b. Ingenuity Pathway Analysis showing the cardiotoxicity report **(B)** and the pathways indicated **(C)** from the differentially expressed genes in non-ischemic heart tissue derived from WT and MPC1$^{CKO}$ heart tissue following I/R injury.

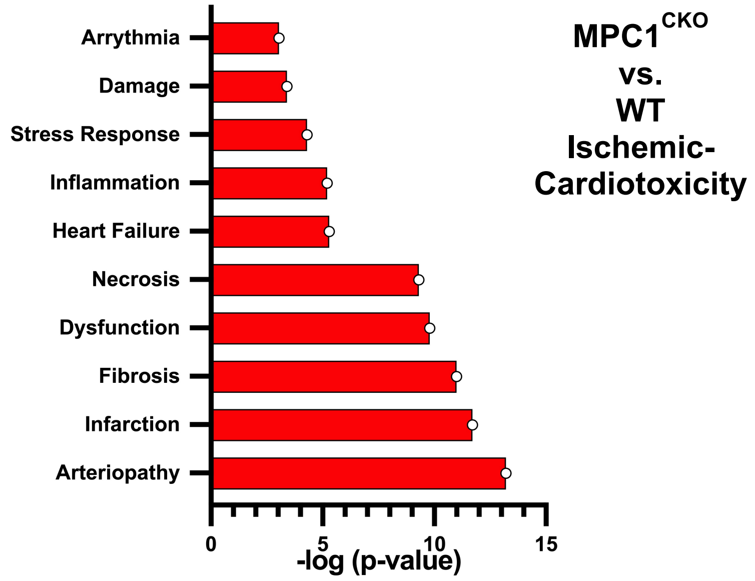


**Supplemental Figure 3:** Cardiotoxicity report from IPA analysis showing that the MPC1$^{CKO}$ genes that were identified in the ischemic myocardium are associated with pathological conditions such as arteriopathy, infarction, and fibrosis.

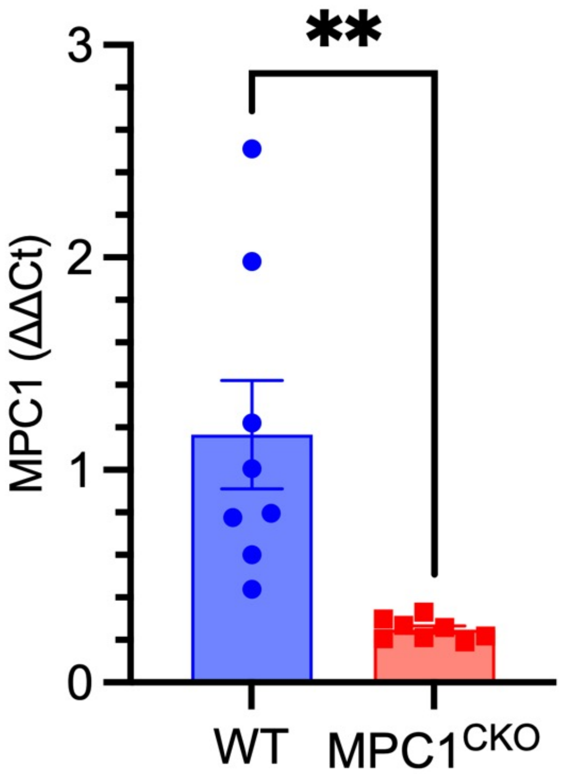


**Supplemental Figure 4:** Quantitative real-time PCR (qRT-PCR) assay for the determination of MPC1 gene expression levels. Expression levels are shown for WT (n=8) and MPC1$^{CKO}$ (n=8) mouse hearts. Error bars denote SEM. **,p-value < 0.01, indicating a significant reduction in MPC1 expression in the MPC1$^{CKO}$ samples compared to WT controls.

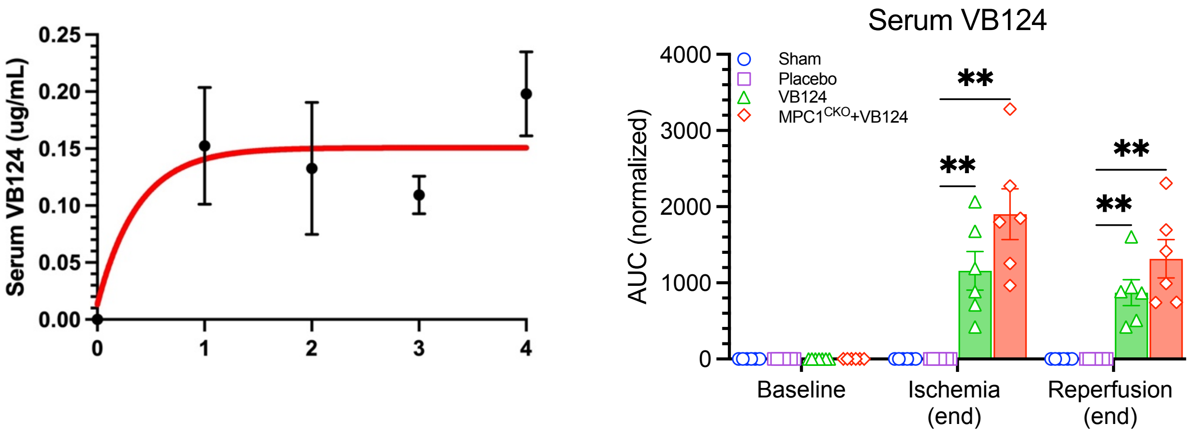


**Supplemental Figure 5:** Serum pharmacokinetics of VB124 post-gavage. **Left)** Serum concentration-time profile of VB124 (30mg/kg) following oral administration (gavage) over a period of 4 hours. The x-axis represents the time post-administration in hours, and the y-axis is the serum concentration of VB124 in micrograms per milliliter (µg/mL). Each point on the curve represents the mean serum concentration of VB124 at the corresponding time point, with error bars indicating SEM from four individual C57BL/6J mice (n=4). The red curve is a “line of best-fit” that models VB124 absorption into the circulation over time. From this data, the 1-hour time-point was selected as our target for reperfusion. **Right)** LC-MS detection of VB124 in sham, placebo, C57Bl/6 mice dosed with VB124, and MPC1$^{CKO}$ mice dosed with VB124.
